# Double training reveals an interval-invariant subsecond temporal structure in the brain

**DOI:** 10.1101/2024.04.30.591981

**Authors:** Shu-Chen Guan, Ying-Zi Xiong, Cong Yu

## Abstract

Subsecond temporal perception is critical for understanding time-varying events. Many studies suggest that subsecond timing is an intrinsic property of neural dynamics, distributed across sensory modalities and brain areas. Furthermore, we hypothesize the existence of a more abstract and conceptual representation of subsecond time, which may guide the temporal processing of distributed mechanisms. However, one major challenge to this hypothesis is that learning in temporal interval discrimination (TID) consistently fails to transfer from trained intervals to untrained intervals. To address this issue, here we examined whether this interval specificity can be removed with double training, a procedure we originally created to eliminate various specificities in visual perceptual learning. Specifically, participants practiced the primary TID task, the learning of which per se was specific to the trained interval (e.g., 100 ms). In addition, they also received exposure to a new interval (e.g., 200 ms) through a secondary and functionally independent tone-frequency discrimination (FD) task. This double training successfully enabled complete transfer of TID learning to the new interval, indicating that training improved an interval-invariant component of temporal interval perception, which supports our general proposal of an abstract and conceptual representation of subsecond time in the brain.

## Introduction

Accurate timing is crucial in perceiving sensory and cognitive events that change dynamically with subsecond precision, like speech and music. Initially, it was believed that subsecond timing relies on a dedicated centralized clock, acting like a pacemaker and accumulator to keep track of time (Creelman, 1962; Treisman, 1963). However, subsequent studies revealed that timing is more likely an intrinsic property of neural dynamics, distributed across sensory modalities and brain areas (Ivry & Schlerf, 2008; Paton & Buonomano, 2018). Various mechanisms contribute to subsecond timing, which for instance include temporal ramps in neuronal firing (Durstewitz, 2003), sequential activation patterns of neuron pools (Hass, Blaschke, Rammsayer, & Herrmann, 2008), the temporal evolution of collective neural network state (Buonomano, 2000), and cortical oscillations at different frequency bands (Matell & Meck, 2004; Allman, Teki, Griffiths, & Meck, 2014). All these mechanisms provide sufficient information for subsecond timing (Hass & Durstewitz, 2016).

Despite the overwhelming evidence in favor of distributed timing mechanisms, including psychophysical evidence such as modality- and duration-specific adaptation (Johnston, Arnold, & Nishida, 2006; Burr, Tozzi, & Morrone, 2007; Heron et al., 2012; Bruno & Cicchini, 2016) and perceptual learning (Lapid, Ulrich, & Rammsayer, 2009; Bratzke, Seifried, & Ulrich, 2012; McGovern, Astle, Clavin, & Newell, 2016), there is also evidence for more general timing mechanisms. For example, a modality-unspecific component of time perception has been suggested by computer simulation (Bratzke & Ulrich, 2019), structural equation modeling of experimental data (Stauffer, Haldemann, Troche, & Rammsayer, 2012), and more direct crossmodal interference of duration judgments (Filippopoulos, Hallworth, Lee, & Wearden, 2013) and EEG measurements (Barne et al., 2018). Furthermore, our recent report reveals that modality-specific perceptual learning of temporal interval discrimination can actually transfer completely across modalities with a double training procedure (Xiong, Guan, & Yu, 2022), providing further evidence for modality-unspecific temporal processing.

We hypothesize that there exists a general, abstract, and conceptual representation of subsecond time in the brain. This hypothesis has been inspired by our general proposal that perceptual learning is the learning of sensory concepts, so that learning can transfer between stimuli with distinct physical appearances and precisions (thresholds), and between sensory modalities (Wang et al., 2016; Xie & Yu, 2019; Xiong, Tang, Zhang, & Yu, 2020; Hu, Wen, Chen, & Yu, 2021). A conceptual representation of subsecond time is consistent with modality-unspecific temporal processing (Stauffer et al., 2012; Filippopoulos et al., 2013; Barne et al., 2018; Bratzke & Ulrich, 2019; Xiong et al., 2022). However, it faces a serious challenge as previous studies have consistently reported no transfer of TID learning from trained to untrained temporal intervals. Wright, Buonomano, Mahncke, and Merzenich (1997) first reported interval specificity in TID perceptual learning, finding that learning a 100-ms interval between two sounds did not transfer to a 50-ms or 200-ms interval. This observation has been replicated in various studies on sensory and motor TID learning (Nagarajan, Blake, Wright, Byl, & Merzenich, 1998; Meegan, Aslin, & Jacobs, 2000; Karmarkar & Buonomano, 2003; Wright, Wilson, & Sabin, 2010), with the exception of Lapid et al. (2009). There is also evidence supporting the presence of duration channels in the brain, each of which is tuned to a narrow range of sub-second time (Heron et al., 2012; Bruno & Cicchini, 2016; Protopapa et al., 2019). Therefore, the discrepancy between the proposed conceptual time representation and observed interval specificity needs to be resolved, and is indeed resolvable.

In this study, we tested the possibility to use double training to enable TID learning transfer to untrained intervals. Double training is a procedure we developed originally for visual perceptual learning research, which can enable location- or orientation-specific learning to transfer to an untrained retinal location or orientation (Xiao et al., 2008; Zhang et al., 2010). In the current context, it comprised two distinct components: The primary training focused on auditory TID training at a specific time interval, and the secondary training was a tone FD task at an untrained transfer interval. Through control experiments, we proved that this secondary task was functionally independent of the primary task and did not significantly affect TID performance on its own. Our hypothesis was that if TID learning improves an abstract and conceptual representation of subsecond timing, the secondary task could activate specific timing mechanisms that respond to the untrained interval. This, in turn, would facilitate functional connections between the training-improved conceptual representation and temporal inputs associated with the untrained interval, thereby enhancing TID performance at the untrained interval.

## Methods

### Participants

The data reported in this research were collected between 2016 and 2021, from 16 participants (10 females, 20.6 ± 2.4 years old) in Experiment 1, 16 participants (11 females, 20.1 ± 2.4 years old) in Experiment 2, and 15 participants (11 females, 19.9±1.8 years old) in Experiment 3. All participants had either normal vision or vision corrected to normal, and normal hearing (pure-tone thresholds ≤ 20 dB hearing level across the frequency range of 0.5–6 kHz). They had no prior experience with psychophysical experiments, and were naïve to the study’s purpose. Each participant provided informed consent prior to data collection. The study was approved by the Peking University Institutional Review Board and followed the ethical guidelines of the World Medical Association (Declaration of Helsinki) throughout the experiments.

### Transparency and Openness

We provided a comprehensive account of how we arrived at our sample size, all exclusions made to the data (if applicable), all manipulations employed, and the measures taken in our study. Upon request, we will provide access to all data, analysis code, and research materials. Data analysis was conducted using the R software (R_Core_Team, 2015). The design and analysis of this study were not pre-registered.

### Sample Size

The decision for our sample size was based on a previous TID learning study by Wright et al. (1997) that utilized similar stimuli (their Figure 4, 100 ms – 1 kHz condition). For power analysis, we used the G*Power software. In our study, the measures of learning and transfer first involved comparing pre- and post-training thresholds in all experiments. The sample size for each group was thus determined using the t tests family for the difference between two dependent measures (matched pairs). To achieve 80% power at a significance level of *p* = .05, and an effect size of Cohen’s *d* = 1.34 in Wright et al. (1997), a sample size of 7 participants would be necessary. Additionally, when comparing the learning effects among the conventional single training, double training, and control groups, a sample size of 7 participants would be sufficient to detect an interaction with a median effect size of 0.4 between the experimental groups and the pre/post-training with 80% power at p = 0.05. To account for potential dropouts among participants, we determined a sample size of 8 for all experiments.

### Apparatus, Stimuli and Procedures

Experiments were conducted in a soundproof anechoic booth. The stimuli were diotic sound generated using a Matlab-based Psychtoolbox software (Pelli, 1997). These stimuli were then presented to the participants using a pair of Sennheiser HD-499 headphones. Two tasks were designed for this study: the temporal interval discrimination (TID) task and the tone frequency discrimination (FD) task.

#### Temporal Interval Discrimination Task

In the TID task, the stimuli consisted of two 15-ms tone pips separated by various intervals. Each tone pip had a 5-ms cosine ramp at both ends and remained constant at 1 kHz and 86 dB sound pressure level. The specific interval length was determined by calculating the difference between the offset of the first stimulus and the onset of the second stimulus. The TID thresholds were assessed using the method of constant stimuli. In each forced-choice trial, a visual fixation point appeared at the center of the computer screen for 300 ms. Then, two pairs of stimuli were presented in random order, with one pair containing a standard interval (SI) and the other pair containing a comparison interval (SI + ΔI). There was a 900-ms inter-interval time gap between the presentation of these two pairs (see Figure 1A for details). The standard interval varied at 100 ms, 200 ms, or 400 ms depending on the experimental condition. Participants were required to press either the left or right arrow key on the computer keyboard to indicate whether the first or the second pair of stimuli had a longer interval. Following each response, a happy or sad cartoon face appeared on the screen, indicating whether the response was correct or incorrect. A blank screen was then shown for a random duration of 500-1000 ms before the start of the next trial. The ΔI was individually adjusted at six levels for each condition, ensuring an adequate range of correct response rates. Before starting the experiment, each observer underwent a quick practice session. This session provided an initial threshold for the participant, enabling the experimenters to define six levels of ΔI. These levels aimed to cover a probability range from 20% to 80%. For most participants, during the pre-training session, the task at 100-ms intervals typically featured a range of six levels varying from ±20% to ±31%, with two logarithmic steps in negative and positive directions, respectively. Daily thresholds were calculated for each participant during training, and the range of six levels was adjusted for the next day. For tasks at 200-ms and 400-ms intervals, which had lower thresholds, the pre-training ranges were set at ±15% to ±22% and ±12.5% to ±18.75%, respectively. Each level was repeated 10 times within a block of 60 trials. Following the training session, we calculated the daily threshold for each participant and adjusted their six-level range based on the previous day’s threshold.

**Figure 1.**
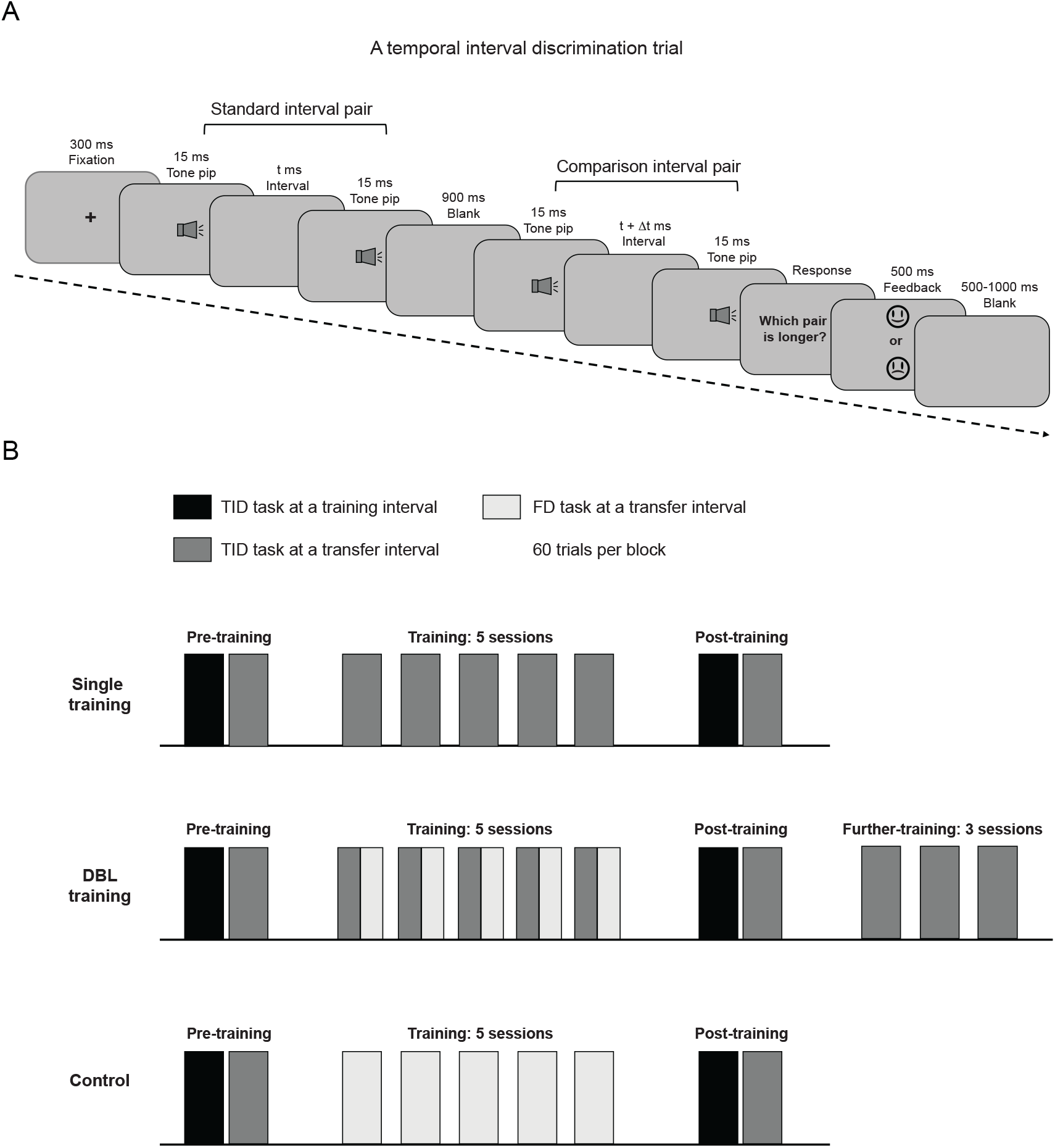
Experimental Task and Design. *Note*. (A) The procedure of a temporal interval discrimination (TID) trial. The standard interval pair included two 15-ms tone pips (1 kHz) separated by a standard interval (SI), whereas the comparison interval pair included the same two-tone pips separated by an SI + ΔI interval. In each trial, the standard and comparison pairs were presented in a random order with a 900-ms time gap. Participants had to determine which pair had a longer interval. For FD trials only conducted in Experiments 2 and 3, the procedure was nearly the same. However, both pairs had an interval identical to the transfer interval (e.g., 200 ms in Experiment 2 and 400 ms in Experiment 3), with one pair having tone pips at a standard frequency (1 kHz), and another pair having tone pips at a higher comparison frequency (1 kHz + Δf). Participants were required to select the pair with a higher frequency. (B) The protocols for three experimental conditions. The pre-training and post-training sessions were identical across all three conditions. These sessions involved TID tasks at the trained and transfer intervals. During the training phase, two single-training groups (Experiment 1) completed five training sessions of the TID task at the training interval. The two double-training groups (Experiments 2 & 3) underwent five training sessions that encompassed both the TID task (at the training interval) and the FD task (at the transfer interval). Lastly, the two control groups (Experiments 2 & 3) completed five training sessions that solely focused on the FD task at the transfer interval. In addition, after the post-training session, the double-training groups underwent three additional sessions of TID training at the transfer interval. This step aimed to evaluate the extent of transfer achieved. Detailed stimulus information were provided in Table S1 in the Supplementary Materials.

For each session, the accuracy data cross the six levels of ΔI were pooled, and a psychometric function was fitted with the equation 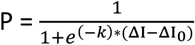, where P represented the rate of reporting the comparison interval being longer at each ΔI, k represented the slope, and ΔI_0_ represented the point of subjective equivalence. A root mean square error (RMSE) value was calculated for each fitted psychometric function as an indicator of goodness of fit. Across all sessions, the RMSEs ranged from 0.01 to 0.11, indicating satisfactory data fitting. The TID threshold was defined as half the interquartile range of the function: Threshold = (ΔI_0.75_-ΔI_0.25_)/2. This threshold represented the ΔI at which participants perceived the comparison interval as shorter than the standard interval in 50% of the trials, and longer than the standard interval in the remaining 50% of trials. Individual pre-training and post-training psychometric functions were provided in Figures S1 -S6 in the Supplementary Materials.

#### Frequency Discrimination Task

In the FD task, the stimuli were identical to those used for the TID task, except that the frequency of the comparison tone pip pair was varied while the temporal interval remained fixed. In each trial, two pairs of tone pips were presented in a random order: one pair at a standard frequency of 1 kHz and the other pair at a higher comparison frequency (1 kHz + Δf). Participants indicated whether the first or second pair of tone pips had a higher frequency by pressing the left or right arrow key. The FD thresholds were assessed with a temporal 2AFC staircase procedure. Initially, the frequency difference (Δf) between the standard and comparison stimuli was set at 50%. This difference decreased by a factor of 2 after every correct response until the first incorrect response occurred. Subsequently, the Δf was adjusted by a factor of 1.414 following a 3-down-1-up staircase rule, aiming for a 79% correct rate. Each staircase terminated after 60 trials. The threshold value was determined as the mean of the last 40 trials.

#### Procedure

In Experiment 1, the participants were randomly assigned to one of two groups with equal numbers. One group of participants (N=8) was trained with a 100-ms interval and assessed for transfer effects at a 200-ms interval. Another group (N=8) was trained with a 200-ms interval and assessed for transfer effects at both 100-ms and 400-ms intervals. Participants completed a pre-training session with TID tasks at both the training and transfer intervals, consisting of 5 blocks for each interval. They then underwent five training sessions, each comprising 16 blocks of the TID task at the training interval, lasting approximately 1.5 hours. A post-training session identical to the pre-training session followed. All sessions were conducted on separate days (Figure 1B), with blocks in each session counterbalanced across intervals. The entire experiment was completed within 7-13 days, with gaps no more than two days between daily sessions.

In Experiment 2, the transfer of TID learning from the 100-ms interval to the 200-ms interval was assessed using a double-training approach. The double-training group (N = 8) underwent a pre-training session, followed by five double-training sessions, a post-training session, and three further- training sessions. The pre- and post-training sessions were the same as those in Experiment 1 (the 100-ms training group). Each double-training session, lasting approximately 2 hours, included both the primary task (TID at a 100-ms interval) and the secondary task (FD at a 200-ms interval), with 10 blocks of each task alternated and counterbalanced. Each further-training session involved 16 blocks of TID task at the 200-ms transfer interval. The control group (N = 8) underwent the same pre- and post-training sessions but solely completed the FD task at the 200-ms interval during five training sessions, with 16 blocks per session. Unlike the double-training group, the control group did not undergo any further training.

In Experiment 3, the transfer of TID learning from the 200-ms interval to the 400-ms interval was assessed using a double-training approach. The double-training group (N = 8) underwent the same protocol as did the double-training group in the Experiment 2, but with difference in the training (200 ms) and transfer (400 ms) intervals. The control group (N = 7) was trained solely on the FD task at the 400-ms interval during five training sessions, following a similar protocol to the control group in Experiment 2.

Table S1 in the supplementary documents summarized the stimulus information for each pre-test, training, and post-test sessions.

### Statistical Analysis

The TID thresholds were initially log-transformed to ensure normal distributions (pre-log-transformation Shapiro-Wilk test: *p* < .01 for TID thresholds at 100-ms, 200-ms, and 400-ms intervals; post-log-transformation Shapiro-Wilk test: *p* = 0.37, 0.10, and 0.57, respectively, for corresponding TID thresholds).

To minimize Type-I error, data from all three experiments were combined into a single linear mixed effects (LME) model to analyze the effects of TID training and transfer. This analysis utilized the “lmer” function from the “lme4” package (Pinheiro & Bates, 2000). The model considered Threshold as the dependent variable and included Group (all six groups across Experiments 1, 2 & 3), Interval (100 ms, 200 ms, and 400 ms), and Test (pre- & post-training) as fixed effects. Individual variations in threshold across Intervals and Tests were modelled as a random effect. We started with a full model in which Interval and Test were each nested within participant. We then constructed reduced model with only Interval or only Test nested within participant. The full model and reduced models were compared using the Likelihood tests to examine which model explained most variations. The best model, in this case the full model, was submitted for analyzing the main effects of the fixed factors. The significance of the fixed effects was assessed by the ANOVA function in the “lmerTest”.

Post-hoc analyses were conducted based on the best fitting model, examining the learning and transfer effects through pair-wise comparisons between pre- and post-training thresholds for each condition in each experiment. Bonferroni correction was applied using the “emmeans” package (Piepho, 2004) during the post-hoc analysis. It is important to note that the absence of significance in the post-hoc analysis does not necessarily suggest the acceptance of the null hypothesis. A Bayesian analysis was thus conducted using JASP to calculate the Bayes factor (BF_10_), which quantifies evidence for the alternative hypothesis (H1) over the null hypothesis (H0) from Bayesian t-tests and ANOVA. A BF_10_ above 1 suggests evidence for H1, a BF_10_ of 1 implies equal support for both hypotheses, and a BF_10_ below 1 favors H0. If requested, all relevant data and code of this study will be provided.

## Results

Since all experiments (1, 2, & 3) were analyzed together using a single linear mixed effects (LME) model (see Statistical Analysis), we began by presenting the overall effects here. The LME results showed significant main effects of Interval (*F*(2, 18) = 32.63, *p* < .001) and Test (*F*(1, 36) = 29.65, *p* < .001), but no significant main effect of Group (*F*(5, 38) = 1.18, *p* = 0.34). Additionally, there were significant interactions between Interval and Test (*F*(2, 30) = 8.69, *p* = .001), as well as among Group, Interval, and Test (*F*(3, 30) = 3.94, *p* = .017). These interactions indicated that the amount of improvement in the posttest was dependent on the task interval and experimental groups. We report experiment-specific effects below.

### Experiment 1: Interval-specificity in TID learning with conventional single training

We first replicated the interval specificity of TID learning in two single-training groups. In the single-training group with a 100-ms TID (N = 8), training led to a significant reduction in TID thresholds at the 100-ms interval. Specifically, the thresholds decreased by 0.29 ± 0.07 log units (*t* = -4.62, *p* < .001, Cohen’s *d* = 1.63, 95% CI [-0.41, -0.16], BF_10_ = 272.94) (red circles in Figure 2A, B). However, training had little impact on TID at the 200-ms interval, reducing the thresholds by merely 0.08 ± 0.04 log units (*t* = - 1.25, *p* = .22, Cohen’s *d* = 0.44, 95% CI [-0.20, 0.05], BF_10_ = 1.48) (green circles in Figure 2A, B), which replicated the well-established interval specificity in TID learning.

**Figure 2.**
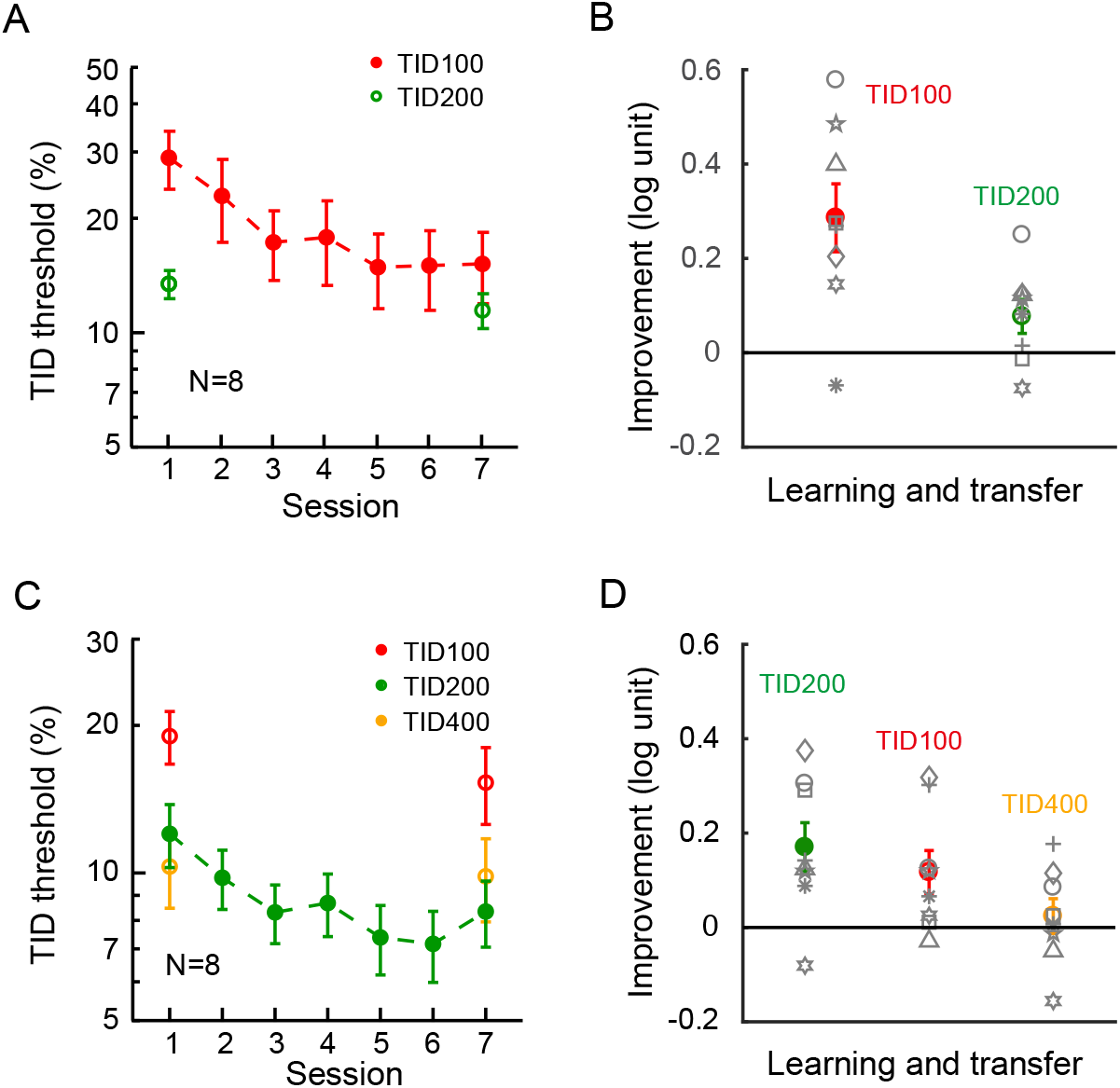
Baselines: Interval Specificity in TID Learning with Conventional Single Training. *Note*. (A) The averaged learning curve for the 100-ms TID task, as well as the pre- and post-training 200-ms TID thresholds. (B) The averaged and individual performance improvements for the trained 100-ms TID task and the untrained 200-ms TID task. (C) The averaged learning curve for the 200-ms TID task, along with the pre- and post-training thresholds for the 100-ms and 400-ms TID tasks. (D) The averaged and individual performance improvements for the trained 200-ms TID, as well as the untrained 100-ms and 400-ms TID tasks. In the plots, the error bars represent ±1 standard error of the mean, and the red, green, and yellow circles correspond to the 100-ms, 200-ms, and 400-ms TID tasks, respectively. In (A) and (C), hollow circles represent untrained tasks that were only performed during Session 1 and Session 7, while solid circles represent the trained tasks that were performed daily.

In the single-training group with a 200-ms TID (N = 8), training significantly reduced TID thresholds at the 200-ms interval by 0.17 ± 0.05 log units (*t* = -2.75, *p* = .008, Cohen’s *d* = 0.97, 95% CI [-0.29, -0.05], BF_10_ = 33.63) (green circles in Figure 2C, D). On the other hand, TID thresholds at the 400-ms interval were barely changed. The thresholds were only reduced by 0.02 ± 0.04 log units (*t* = -0.39, *p* = 0.70, Cohen’s *d* = 0.14, 95% CI [-0.15, 0.10], BF_10_ = 0.42) (yellow circles in Figure 2C, D), which again demonstrated the interval specificity. Interestingly, training also resulted in a reduction in TID threshold at the 100-ms interval by 0.12 ± 0.05 log units (*t* = -1.90, *p* = .063, Cohen’s *d* = 0.67, 95% CI [-0.24, 0.01], BF_10_ = 7.00) (red circles in Figure 2C, D). This reduction was replicated using non-log-transformed raw thresholds (threshold reduction = 3.73 ms, t = -1.87, p = .066, Cohen’s d = 0.66). Therefore, the learning from the 200-ms TID task partially transferred to the 100-ms interval, which was approximately 40% of the direct training effect observed with the 100-ms TID task (red circles in Figure 2B).

### Experiment 2: TID learning transfer from a 100-ms trained interval to a 200-ms interval with double training

Consistent with our hypothesis, we found a significant transfer effect from 100-ms to 200-ms after double training. Double training resulted in a significant reduction in 100-ms TID thresholds by 0.27 ± 0.06 log units (*t* = -4.30, *p* < .001, Cohen’s *d* = 1.52, 95% CI [-0.39, -0.14], BF_10_ = 956.31) (red circles in Figure 3A, C). Notably, 200-ms TID thresholds also exhibited a reduction of 0.18 ± 0.03 log units (*t* = - 2.91, *p* = .005, Cohen’s *d* = 1.02, 95% CI [-0.30, -0.06], BF_10_ = 120.19) (TID 200 in Figure 3C). This reduction was not significantly different from the threshold reduction observed during direct 200-ms TID training in Figure 2D and Figure 4C (green circles) (*F*(2,21) = 0.43, *p* = .66, BF_10_ = 0.43).

**Figure 3.**
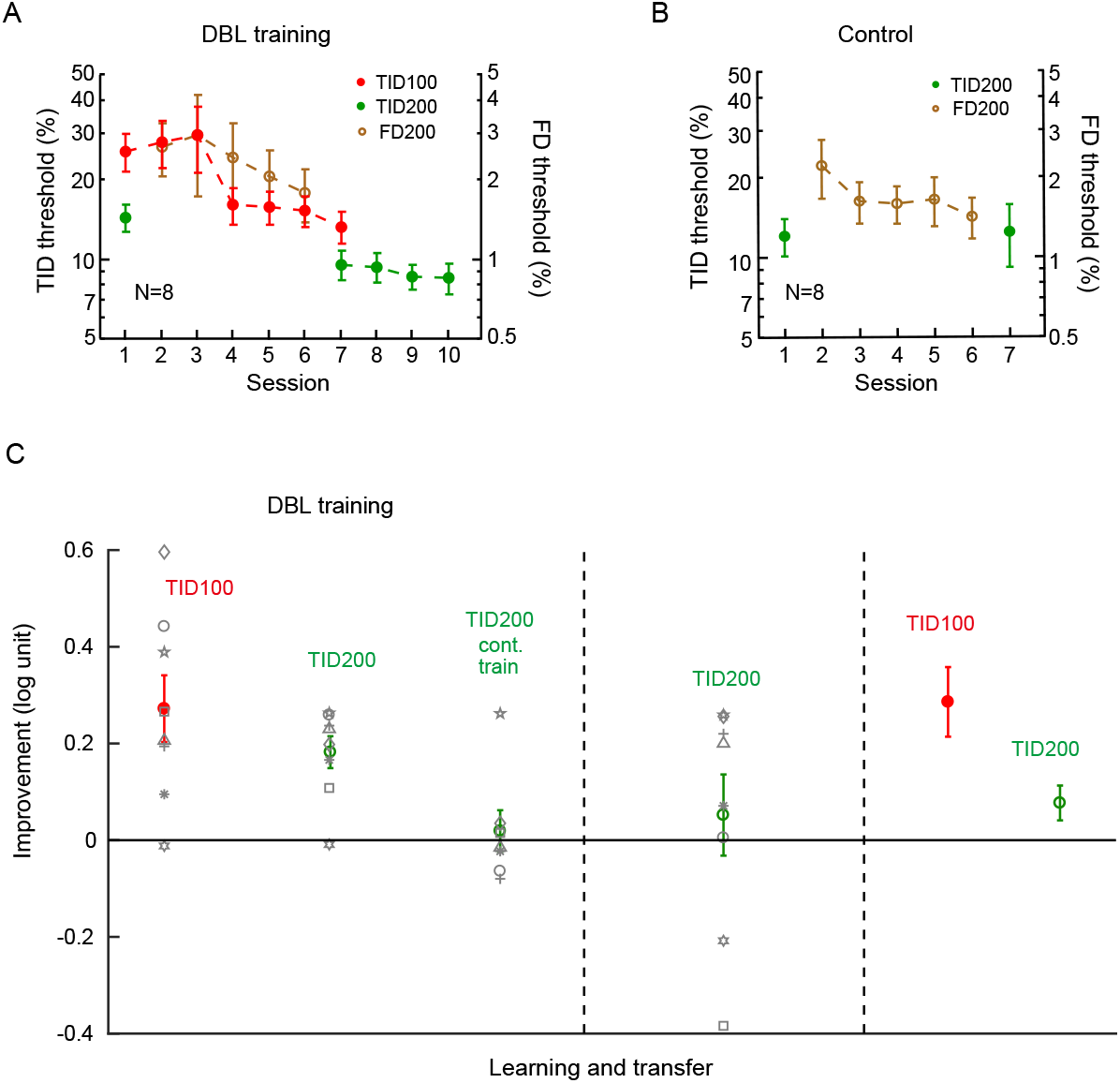
Transfer of TID Learning from 100 ms to 200 ms with Double Training. *Note*. (A) The averaged learning progress of double training for the 100-ms TID task (Sessions 1-7, the curve with red circles) and the 200-ms frequency discrimination (FD) task (Sessions 2-6, the curve with brown circles). The evaluation of the 200-ms TID task was conducted during the initial and final double training sessions (Sessions 1 and 7, green circles). Additionally, the participants practiced the 200-ms TID task for additional 3 sessions (Sessions 8-10). (B) The effect of exclusive 200-ms FD training (the curve with brown circles) on 200-ms TID thresholds (green circles), serving as a control measure. (C) The averaged improvements and individual progressions resulting from double training (100-ms TID and 200-ms TID) and the control condition (200-ms TID). To facilitate comparison, the earlier single-training data for the 100-ms TID task in Figure 2B were re-plotted here). The error bars represent ±1 standard error of the mean.

**Figure 4.**
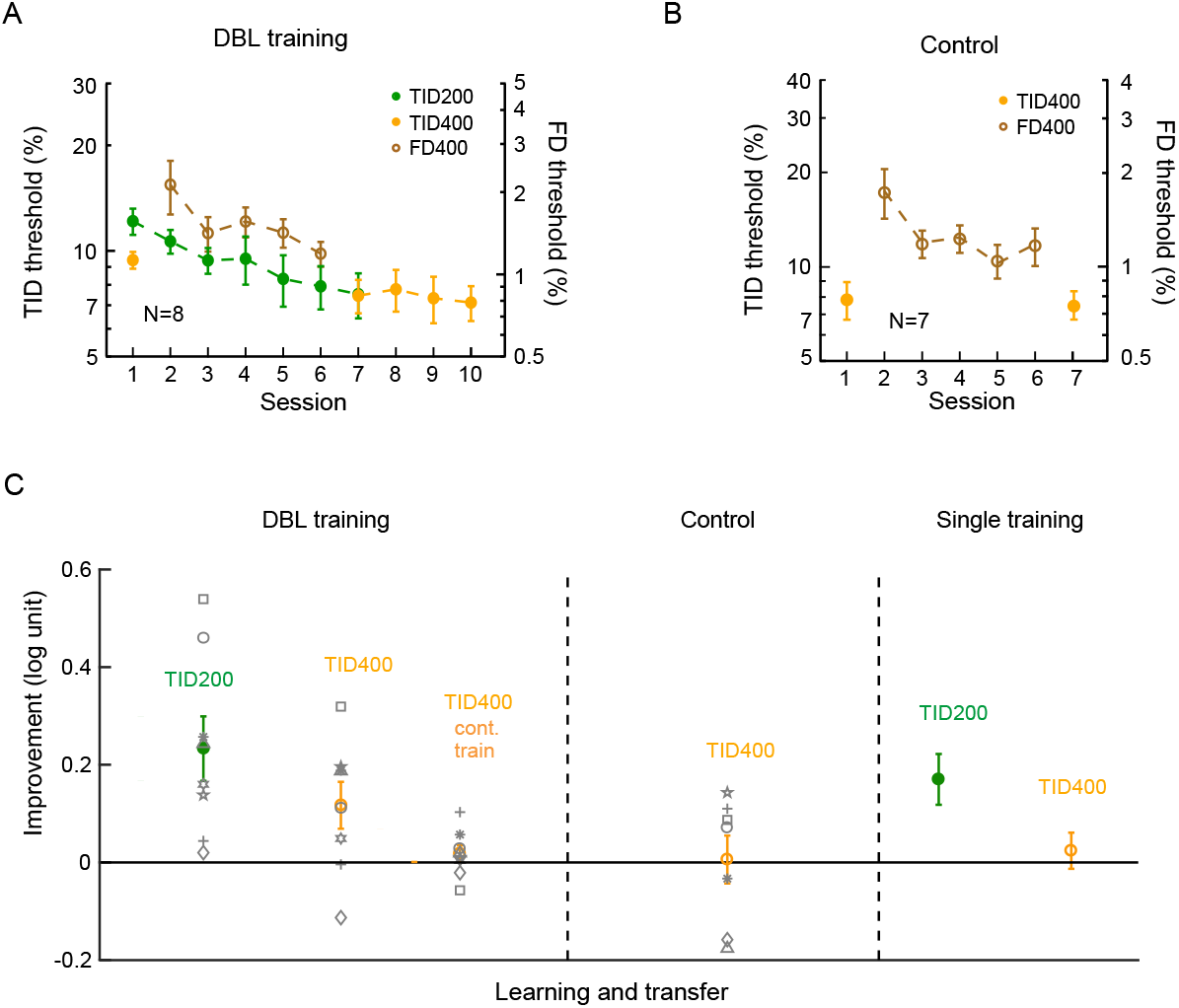
Transfer of TID Learning from 200 ms to 400 ms with Double Training. *Note*. (A) The averaged learning curves for double training, which included 200-ms TID (Sessions 1-7, curve with green circles) and 400-ms frequency discrimination (FD) (Sessions 2-6, curve with brown circles). The 400-ms TID thresholds were assessed in both pre- and post-double training sessions (Sessions 1 & 7, yellow circles), and additional practiced for 400-ms TID occurred over 3 sessions (Sessions 8-10, curve with yellow circles). (B) The impact of 400-ms FD training (curve with brown circles) alone on 400-ms TID thresholds (yellow circles), which served as a control measure. (C) The averaged and individual improvements resulting from double training (200-ms TID & 400-ms TID, left side), as well as the control condition (400-ms FD, middle). The previous data on 200-ms TID single-training were replotted from Figure 2D for comparison. The error bars indicate ±1 standard error of the mean.

To explore whether the transfer effect had reached its maximum, a subgroup of seven participants engaged in an additional three sessions of practicing 200-ms TID, consisting of 16 blocks per session. However, these additional sessions failed to further enhance TID performance at this interval (TID 200 cont.train in Figure 3C) (by 0.001 ± 0.06 log units; t(6) = 0.01, *p* = .99, Cohen’s d = 0.003, BF_10_ = 0.35). These results collectively suggest that the performance of 200-ms TID reached its peak after a combination of both 100-ms TID training and 200-ms tone frequency training in the double training paradigm, despite the fact that it was unaffected by 100-ms TID training alone (green circles in Figure 2A, B).

Since the double training paradigm introduced a new task, it was important to exclude the possibility that the new task per se led to the TID improvement at the 200-ms interval. To account for this alternative possibility, a separate control group (N = 8) only practiced tone FD at the 200-ms interval between the pre- and post-training sessions. The FD training resulted in a significant improvement in 200-ms tone FD, indicated by an improvement of 0.18 ± 0.07 log units (brown circles in Figure 3B). However, this practice did not yield a significant impact on 200-ms TID thresholds (-0.01 ± 0.11 log units; t = -0.20, p = .84, Cohen’s d = 0.07, 95% CI [-0.14, 0.11], BF_10_ = 0.34 (green circles in Figure 3B, C). The outcomes of double training and the control group imply that the combination of 100-ms TID training and 200-ms tone frequency training actuated complete transfer of TID learning from a 100-ms interval to a 200-ms interval, despite the absence of significant transfer in the single-training condition.

### Experiment 3: TID learning transfer from a 200-ms trained interval to a 400-ms interval with double training

We further validated the double training effect by showing a transfer effect from 200-ms TID to 400-ms TID. Double training resulted in a significant change of 200-ms TID, with an improvement of 0.23 ± 0.07 log units (*t* = -3.76, *p* < .001, Cohen’s *d* = 1.33, 95% CI [-0.36, -0.11], BF_10_ = 52.29) (green circles in Figure 4A, C). Similar to the earlier double training experiment, the untrained 400-ms TID task also displayed improvement, with a reduction of TID thresholds by 0.12 ± 0.05 log units. This transfer effect was marginally significant with a medium effect size and a large Bayes factor (*t* = -1.88, *p* = .065, Cohen’s *d* = 0.67, 95% CI [-0.24, 0.01], BF_10_ = 3.24) (yellow circles in Figure 4A, C).

To assess the completeness of the transfer effect, all participants underwent three additional practice sessions for the 400-ms TID, with each session containing 16 blocks of trials. Despite these additional sessions, we observed no significant improvement in TID performance for the 400-ms interval, with the threshold reduced by 0.02 ± 0.02 log units (*t*(7) = - 0.24, p = .82, Cohen’s *d* = 0.08, BF_10_ = 0.35) (TID 400 cont.train in Figure 4C). The outcomes indicate that the learning from 200-ms TID effectively transferred to and optimized the performance of the 400-ms TID following the double training sessions.

Again, a control experiment was also conducted to test the alternative possibility that the secondary task per se may lead to TID improvement at the 400-ms interval. The control group (N = 7) underwent the tone FD training at the 400-ms interval, demonstrated an improvement in the FD performance by 0.15 ± 0.05 log units (brown circles in Figure 4B), but a minimal change of the 400-ms TID thresholds by 0.01 ± 0.05 log units (*t* = -0.10, *p* = .92, Cohen’s *d* = 0.26, 95% CI [-0.14, 0.13], BF_10_ = 0.36 (yellow circles in Figure 4B, C). This implies that the improvement observed in the 400-ms TID improvement was indeed due to double training and was not the result of the 400-ms FD training alone. The pre-training 400-ms TID threshold (yellow circle of Session 1 in Figure 4B) for the control group appeared to be lower than that of the double training group (yellow circle of Session 1 in Figure 4B). This difference was mainly attributable to one participant who had an exceptionally low pre-training threshold at 4.1%. Upon excluding this participant’s data from the analysis, the statistical conclusions remained consistent.

## Discussion

In this study, we first replicated the specificity of TID learning, observing that training at 100-ms and 200-ms intervals did not transfer to untrained 200-ms and 400-ms transfer intervals, respectively (Experiment 1). However, a double training paradigm, in which the primary TID task at the trained interval was combined with a secondary FD task at the transfer interval, led to complete transfer of TID learning (Experiments 2 & 3). Furthermore, control experiments showed that the secondary FD learning task per se could not explain the transfer effect (Experiments 2 & 3).

Time interval information after initial processing by distributed mechanisms such as hypothesized duration channels (Heron et al., 2012; Bruno & Cicchini, 2016; Protopapa et al., 2019) requires subsequent readout by more centralized decision units (Bueti & Buonomano, 2014; Paton & Buonomano, 2018). According to various reweighting theories of perceptual learning (e.g., Dosher & Lu, 1998), training in TID would be expected to enhance the readout of time information by assigning more weight to temporal inputs that best match a stimulus template representing the trained interval. Here the stimulus template is supposedly interval specific, thus predicting interval specificity in TID learning. However, the observations of cross-interval transfer of TID learning in the current study suggest a more general interval-invariant time representation, which is probably a higher-level process than the rigid templates representing specific intervals.

Specificities in perceptual learning have been used to infer the neural mechanisms underlying learning (Sagi, 2011; Watanabe & Sasaki, 2015). In time perception, the interval and modality specificities in TID learning were also interpreted as evidence against a centralized clock and for distributed mechanisms (Bueti & Buonomano, 2014). However, double training results suggest that such specificities likely arise from particular single-training procedures, rather than being inherent properties of temporal learning. One possible cause for learning specificity might be various degrees of overfitting during training (Mollon & Danilova, 1996; Sagi, 2011). According to this account, the participants during training may learn to attend to various peculiarities that are associated with the training condition but are not necessarily relevant to the trained task. Because of overfitting, learning cannot transfer to new conditions where the same peculiarities may not exist. Similarly, we propose that as training requires full attention to the trained condition, it may suppress or ignore untrained conditions that are neither attended no stimulated (Xiao et al., 2008; Xiong, Zhang, & Yu, 2016). As a result, high-level conceptual and transferrable learning cannot functionally connect to sensory inputs from transfer conditions, such as time inputs originating from a different modality or representing a different interval, to enable learning transfer (e.g., Xiao et al., 2008; Zhang et al., 2010; Xiong et al., 2016; Xiong et al., 2022).

Double training enables learning transfer within the same task, such as from one orientation to another orientation or from one retinal location to another retinal location in an orientation or contrast discrimination task, but it fails to make learning transfer across tasks, such as from contrast discrimination to orientation discrimination and vice visa (Cong, Wang, Yu, & Zhang, 2016). These results provide further constraint that learning is specific to a sensory concept, such as an orientation or contrast concept. Following the discovery of the double-training effect, other experimentally introduced factors, such as attention (Donovan, Szpiro, & Carrasco, 2015; Xiong et al., 2016; Donovan & Carrasco, 2018), stimulus variability (Xie & Yu, 2020; Manenti, Dizaji, & Schwiedrzik, 2023), and even subconscious stimulation (Xiong et al., 2016), have also been found effective to actualize learning transfer. While stimulus variability prevents overfitting, the secondary task in double training may override overfitting by informing the brain to ignore task-irrelevant peculiarities learned in the primary training task. Neural network-wise, the secondary task, as well as top-down attention and bottom-up stimulation, can activate neurons responding to the transfer condition, so that functional connections between high-level conceptual learning and new sensory inputs can be established or strengthened to achieve learning transfer (Xiao et al., 2008; Xiong et al., 2016). Such functional connections have been suggested by ERP evidence that significant occipital P1-N1 changes are associated with learning transfer to a transfer location, but are absent when learning fails to transfer (Zhang, Cong, Song, & Yu, 2013).

Is it possible that double-training enabled learning transfer is actually a result of joint activations of a certain mechanism by the primary and secondary tasks, even if neither task can enable the learning transfer on its own? Previously we have discovered an order effect of double training. That is, perceptual learning of visual orientation discrimination and auditory tone frequency discrimination only transfers when the secondary training is conducted either simultaneously with the primary training or at a later time, but not ahead of it (Zhang et al., 2010; Xiong et al., 2020). These findings may be inconsistent with a jointly activated mechanism to underlie double training, unless the mechanism also requires some specific order of activations. We rather interpret the order effect as an indication that perceptual learning of the primary task needs to take place before it transfers (Zhang et al., 2010; Xiong et al., 2020).

The thresholds for 100-ms TID were about twice as high as those for 200-ms TID (Figure 2A), suggesting less precision and higher uncertainty. Notably, even the post-training 100-ms TID thresholds remained higher than the pre-training 200-ms TID thresholds. Therefore, learning with a less precise and more uncertain 100-ms TID task cannot be directly mapped to 200-ms TID. Similarly, due to these precision or uncertainty differences, neither enhanced attention to nor improved memory of the trained temporal intervals resulting from TID training can be directly responsible for the learning transfer from 100-ms to 200-ms. Furthermore, the improvement of this temporal perception is likely task-specific, as we did not observe an improvement of TID task after FD training (Figures 3B, 4B), which excludes the possibility that TID learning transfer is caused by training-improved general decision-making capability, as well as general auditory attention (both are auditory tasks). Instead, the cross-interval learning transfer suggests that training enhances some fundamental knowledge of temporal interval information. This interpretation is consistent with, and thus offers further support to, our proposition that TID training improves an abstract and conceptual representation of subsecond time.

Although the interval and modality specificities of TID learning have been used to dispute a centralized clock, double-training enabled TID learning transfer across intervals and modalities may not necessarily support a dedicated centralized clock. While a centralized clock is assumed to measure time, a conceptual representation of subsecond time lacks the ability to do so since it only involves abstracted time information. Therefore, time needs to be measured by specific timing mechanisms. Furthermore, training of a TID task at higher thresholds (e.g., 100-ms TID) would have no impact on the machinery of the hypothesized centralized clock that is capable of handling more precise timing tasks (e.g., TID at 200 ms), not supporting the idea of a centralized clock either.

Bueti et al. (2012) found that visual cortices may directly contribute to the representation of time in vision modality. In contrast, the insula appears to be involved in “high-level” processing of temporal information that is modality unspecific (Bueti, Lasaponara, Cercignani, & Macaluso, 2012). The detailed brain mechanisms specific to the transfer of TID learning across intervals may be uncovered through future imaging studies. These mechanisms would involve cortical or subcortical structures with an improved ability to decode time interval information after training and effective connections from these structures to sensory areas (e.g., V1 or A1) that facilitate learning transfer.

## Conclusion

In this study, we successfully demonstrated the complete transfer of TID learning from trained to untrained new intervals using a double training approach, even when the same learning was initially interval-specific with conventional (single) training. These findings provide strong evidence for the existence of an interval invariant representation of subsecond time and its precision being enhanced through learning. Prior evidence has already suggested that subsecond time perception is not constrained by sensory modalities (Stauffer et al., 2012; Filippopoulos et al., 2013; Barne et al., 2018; Xiong et al., 2022). However, being general across these non-temporal features is necessary but not sufficient for a general representation of subsecond time, as training-improved precision of this representation shall apply to different time intervals, which would predict cross-interval transfer of temporal learning. This prediction is finally proved by our current findings.

## Constraints on Generality

Our participants were convenient samples of Chinese college students with normal hearing and learning abilities from a diverse background of majors. We expect our findings to generalize to a broad population of adults with normal hearing and learning abilities. However, the learning and transfer effects we observed in this study may not be replicated in subjects with hearing loss, learning disabilities, or cognitive decline. We determined the numbers and durations of the training sessions based on our prior experience in visual and auditory perceptual learning. However, individuals with hearing loss or who are older in age may require longer training courses and exhibit smaller learning and transfer effects. We recommend that the best practice to examine whether a transfer effect is complete is to conduct continuous training to observe whether further improvement can be achieved. We have no reason to believe that the results depend on other characteristics of the participants, materials, or context.

